# Altered circadian behavior and light sensing in mouse models of Alzheimer’s disease

**DOI:** 10.1101/2023.05.02.539086

**Authors:** Thaddeus K. Weigel, Cherry L. Guo, Ali D. Güler, Heather A. Ferris

## Abstract

Circadian symptoms have long been observed in Alzheimer’s disease (AD) and often appear before cognitive symptoms, but the mechanisms underlying circadian alterations in AD are poorly understood. We studied circadian re-entrainment in AD model mice using a “jet lag” paradigm, observing their behavior on a running wheel after a six hour advance in the light:dark cycle. Female 3xTg mice, which carry mutations producing progressive amyloid beta and tau pathology, re-entrained following jet lag more rapidly than age-matched wild type controls at both 8 and 13 months of age. This re-entrainment phenotype has not been previously reported in a murine AD model. Because microglia are activated in AD and in AD models, and inflammation can affect circadian rhythms, we hypothesized that microglia contribute to this re-entrainment phenotype. To test this, we used the colony stimulating factor 1 receptor (CSF1R) inhibitor PLX3397, which rapidly depletes microglia from the brain. Microglia depletion did not alter re-entrainment in either wild type or 3xTg mice, demonstrating that microglia activation is not acutely responsible for the re-entrainment phenotype. To test whether mutant tau pathology is necessary for this behavioral phenotype, we repeated the jet lag behavioral test with the 5xFAD mouse model, which develops amyloid plaques, but not neurofibrillary tangles. As with 3xTg mice, 7-month-old female 5xFAD mice re-entrained more rapidly than controls, demonstrating that mutant tau is not necessary for the re-entrainment phenotype. Because AD pathology affects the retina, we tested whether differences in light sensing may contribute to altered entrainment behavior. 3xTg mice demonstrated heightened negative masking, an SCN-independent circadian behavior measuring responses to different levels of light, and re-entrained dramatically faster than WT mice in a jet lag experiment performed in dim light. 3xTg mice show a heightened sensitivity to light as a circadian cue that may contribute to accelerated photic re-entrainment. Together, these experiments demonstrate novel circadian behavioral phenotypes with heightened responses to photic cues in AD model mice which are not dependent on tauopathy or microglia.

## Introduction

Altered circadian rhythms are a common symptom of Alzheimer’s disease (AD). These alterations appear early in the disease, before hallmarks such as memory impairment, amyloid-β (Aβ) plaques, and neurofibrillary tangles (Musiek et al., 2015). AD circadian symptoms include sleep disruptions and a greater severity of behavioral symptoms later in the day, known as sundowning. Circadian disruptions are also observed at the molecular level, with alterations in circadian clock gene expression in the brains of AD patients (Cermakian et al., 2011). These circadian alterations are particularly interesting because they may play a role in disease progression: sleep can facilitate Aβ clearance from the brain (Shokri-Kojori et al., 2018; Xie et al., 2013) and poor sleep quality in adulthood is a risk factor for AD later in life (Sabia et al., 2021). Additionally, sleep disruptions caused by altered circadian rhythms significantly increase the difficulty of caring for AD patients (Kang et al., 2009; Musiek et al., 2015). Thus understanding the mechanisms of circadian disruption in AD could have both important preventative and therapeutic potential.

Many circadian phenotypes seen in humans with AD are recapitulated in mouse models of AD. Certain AD models demonstrate changes to the free running period (the intrinsic period of an animal’s circadian behavior when kept in constant darkness) and activity in light or dark phases (Sheehan & Musiek, 2020). AD model mice also score better in anxiety tests earlier in their active period compared to later (Bedrosian et al., 2011), a phenotype reminiscent of sundowning in AD patients. Circadian alterations are recapitulated at the molecular level as well, with changes to the amplitude and phase of rhythmic clock gene expression in some AD models including 3xTg (Bellanti et al., 2017) and 5xFAD (Song et al., 2015) mice.

Other facets of circadian rhythms have been less well studied in AD models. Entrainment is the process of synchronizing the biological circadian clock with the daily rhythm of the environment. In this study we tested circadian behavior in models of AD using a “jet lag” protocol. We found that female 3xTg mice re-entrain more rapidly than wild type (WT) controls. We then examined neuroinflammation, amyloid pathology, and changes to light sensing as possible contributors to this altered circadian behavior.

## Results

### 3xTg mice have accelerated circadian re-entrainment

To test the re-entrainment behavior of AD model mice, we first studied female 3xTg mice. The 3xTg mouse model of AD carries pathogenic mutations in amyloid precursor protein (APP), presenilin 1 (PS1), and human tau (MAPT), resulting in progressive accumulation in the brain of Aβ plaques and neurofibrillary tangles. Sex-specific circadian behavioral alterations have previously been observed in 3xTg mice (Sterniczuk et al., 2010), and female 3xTg mice have a more rapid progression of AD pathology than males (Dennison et al., 2021). In a photic phase shift experiment, which measures the shifting of circadian behavior caused by one pulse of light during the dark phase, female 3xTg mice showed a trend towards greater phase shifting while males did not (Sterniczuk et al., 2010). We examined re-entrainment in these mice using a shifted light-dark (LD) cycle, simulating travel across 6 time zones and subsequent “jet lag”. This behavior is not altered in male 3xTg mice (González-Luna et al., 2021), but female mice, which have more severe AD pathology than males, have not been studied in this paradigm. We allowed female 8-month-old 3xTg and B6129SF2/J wild type (WT) control mice to entrain to a 12:12 L:D light cycle and monitored their behavior on a running wheel. At this age, female 3xTg mice have only mild Aβ and tau pathology (Fig. 1A, upper panels). Plaques and phosphorylated tau are not observed in the SCN (Supplementary Fig. 1A). After full entrainment and habituation to the running wheel, the LD cycle was advanced by 6h (Fig. 1B). The onset of nightly running was measured in the days following the light cycle shift. 3xTg activity onset was significantly earlier than WT following the light cycle shift on day 2 (p<.01) after the shift (Fig. 1C), demonstrating more rapid re-entrainment. We calculated the number of days each mouse took to complete half of the re-entrainment, the 50% phase shift (PS_50_), and found that 3xTg mice reached their PS_50_ in on average 1.07 fewer days (p<.006) than WT (Fig. 1D). We examined free-running period when kept in 24h darkness (Fig. 1E) and preference for running during the dark phase (Fig. 1F) and found no difference between 3xTg and WT in these other aspects of circadian behavior. Total running was not affected by genotype (Fig. 1G), suggesting that the wheel running re-entrainment phenotype is not influenced by the hyperactivity or perseverative behavior sometimes observed in AD mouse models.

**Figure 1:**
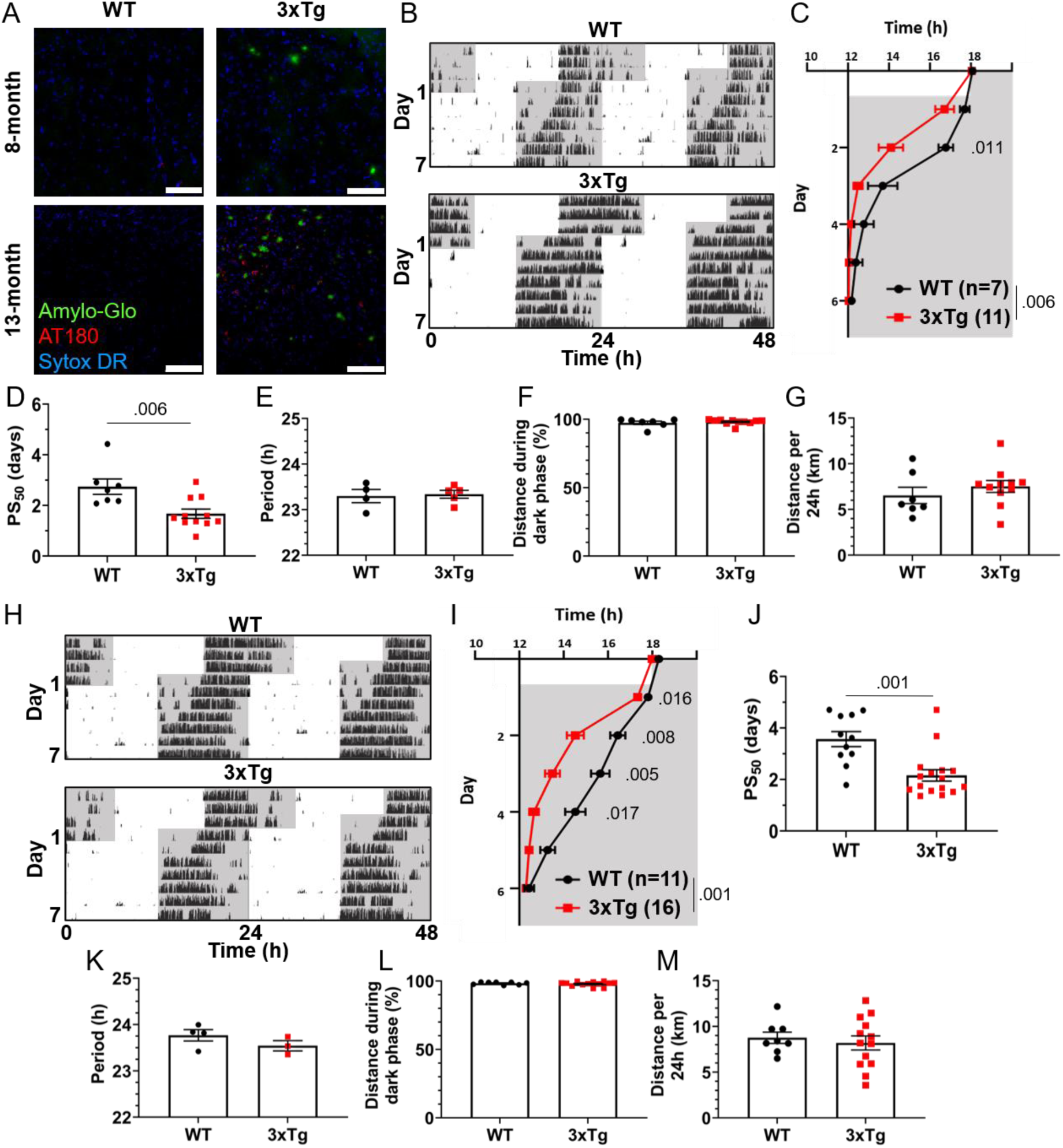
Altered circadian re-entrainment in 3xTg mice. (A) Representative images from the hippocampus of 8-month-old (mo) and 13mo B6129SF2/J wild type (WT) and 3xTg mice. Aβ plaques are stained with Amylo-Glo (green), phosphorylated tau is stained with AT180 (red), and nuclei are stained with Sytox-DR. Scale bars =100um. (B) Representative double-plotted actograms of 8mo WT and 3xTg mice subjected to a 6h phase advance. Light and dark phases of the LD cycle are represented by white and grey background, respectively. (C) Group analysis of activity onset in 8mo mice, with grey representing darkness as in (B). Mixed model with Sidak post hoc comparison, n=7-11. (D) Time to 50% of total phase shift (PS_50_) in 8mo mice from (C), n=7-11. (E) Free-running period (averaged over 7 days) in 8mo mice maintained in constant darkness. (F) Percent of running performed during the dark phase and (G) total distance run in 24 hours (averaged over two 24h periods) in 8mo mice, n=7-11. (H-M) Same as (B-G) but in 13mo mice. n=11-16 in (I-J), 3-4 in (K), 8-13 in (L-M). All analyses are two tailed Student’s t-tests unless otherwise noted. All data plotted as mean ± SEM.

To test whether this phenotype becomes more severe as AD pathology progresses with aging, we repeated this experiment using 13-month-old female 3xTg and WT mice (Fig. 1A, bottom panels; Supplementary Fig. 1B). Behavior onset after the light cycle shift was significantly earlier in 13-month 3xTg than WT mice on days 1 (p<.016), 2 (p<.008), 3 (p<.005), and 4 (p<.017) after the shift (Fig. 1I). Mean PS_50_ at 13 months was 1.41 days faster (p<.001) in 3xTg than WT (Fig. 1J). Free running period and preference for running in the dark phase were not affected by genotype (Fig. 1K-L). Total running was again not affected by genotype (Fig. 1M). These results show that 3xTg mice re-entrain more rapidly in a jet lag paradigm and that this phenotype becomes more severe with aging.

3xTg mice exhibit a metabolic phenotype resulting in greater body weight than B6129SF2/J controls (Robison et al., 2020), and lost more weight during their time with running wheel access, but neither greater body weight nor greater weight loss correlated with more rapid re-entrainment (Supplementary Fig. 2A-F). These metabolic differences are thus not likely to contribute to the observed re-entrainment phenotype.

### Microglia depletion does not alter circadian re-entrainment in 3xTg mice

As neuronal loss does not appear in AD until long after the development of circadian symptoms, we sought other possible mechanisms underlying this behavioral phenotype. Microglia are heavily implicated in AD pathophysiology and microglia activation can contribute to disease progression. In the 3xTg model, the brain has elevated levels of microglia-produced pro-inflammatory cytokines (Park et al., 2021) and microglia activation and proliferation can be observed before the development of Aβ plaques (Janelsins et al., 2005). We observe activated microglia in 3xTg mice at 13mo, where they cluster around Aβ plaques and display a more amoeboid morphology (Fig. 2A). Microglia depletion in AD models decreases neuroinflammatory signaling without acutely altering amyloid and tau pathology and in some studies can partially restore memory deficits (Spangenberg et al., 2016). We hypothesized that activated microglia and neuroinflammation could contribute to the circadian re-entrainment phenotype observed in 3xTg mice and microglia depletion would rescue the re-entrainment phenotype. We used the colony stimulating factor 1 receptor (CSF1R) antagonist Plexxikon 3397 (PLX) to rapidly deplete microglia from the brain. Following re-entrainment to the shifted light cycle in Fig. 1, 13-month 3xTg or WT mice were switched to control or PLX chow (600mg/kg) for 7 days to deplete microglia (Fig. 2B). PLX treatment effectively depleted microglia from the brain in WT and 3xTg mice, reducing the number of microglia in the ventromedial hypothalamus, the region containing the SCN, by >98% (Fig. 2C-D). After 7 days of PLX or control treatment, light cycles were advanced by 6h and running wheel behavior monitored (Fig. 2E). PLX treatment did not rescue the more rapid re-entrainment in 3xTg mice, and there was no significant effect of treatment on time of running onset (Fig. 2F). As in Figure 1, behavior onsets were earlier in 3xTg than WT mice, with a significant effect of genotype on time of running onset (p<.001). PLX treatment did not significantly alter PS_50_ in 3xTg mice (Fig. 2G). There was a nonsignificant trend towards lower PS_50_ in WT control vs. PLX-treated mice (p<.415). There was no significant effect of treatment on PS_50_ (p<.316), but the effect of genotype on PS_50_ was again significant (p<.001). Thus, acute microglia depletion did not rescue the re-entrainment phenotype observed in 3xTg mice. PLX treatment also did not alter other running behaviors measured, with no significant effect of genotype or treatment found in percent running during the dark phase or total distance traveled (Fig. 2H-I). These data demonstrate that re-entrainment remains altered in microglia-depleted 3xTg mice, suggesting that microglia and neuroinflammation are not acutely responsible for this circadian phenotype.

**Figure 2:**
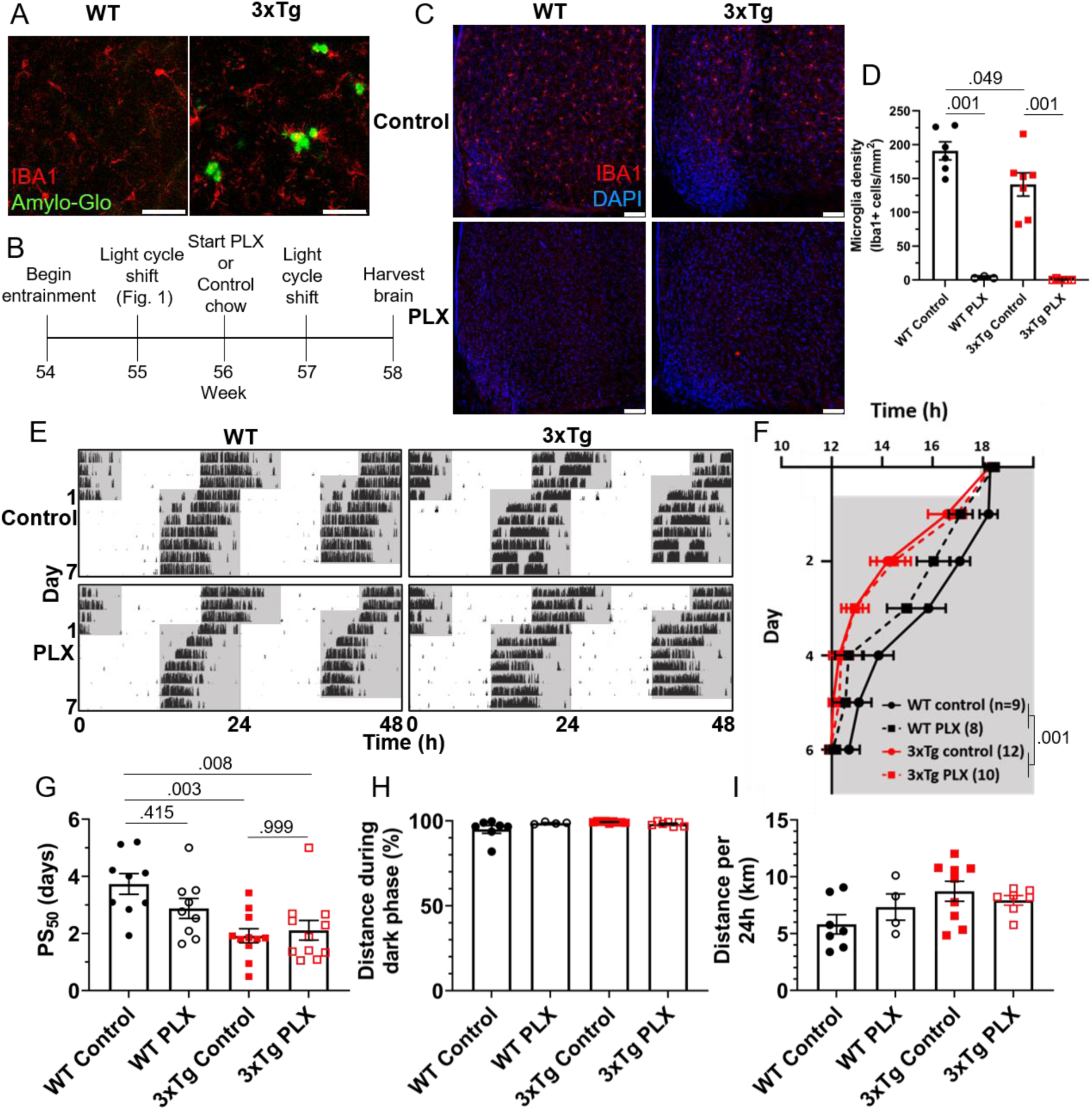
Microglia depletion does not rescue circadian re-entrainment phenotype in 3xTg mice. (A) Representative images of hippocampal microglia morphology in 13mo 3xTg and B6129SF2/J WT mice. Aβ plaques are stained with Amylo-Glo (green) and microglia are marked by staining for IBA1 (red). Scale bars =50um. (B) Timeline of experiment. After completing the light cycle shift experiment in Fig. 1B-D, 13mo WT and 3xTg mice were fed Plexxikon 3397 (PLX) chow to deplete microglia or control chow for 7 days before beginning light cycle shift. (C) Representative images of microglia depletion in the SCN and surrounding region in WT and 3xTg mice fed control or PLX diets. Microglia labeled by staining for IBA1 (red). Scale bars =100um. (D) Quantification of microglia in the ventromedial hypothalamus, the region containing the SCN, in WT and 3xTg mice fed control or PLX diets. 2-way ANOVA. (E) Representative double-plotted actograms of 13mo wild type (WT) and 3xTg mice, treated with control or PLX chow, subjected to a 6h phase advance. Light and dark phases of the LD cycle are represented by white and grey background, respectively. (F) Group analysis of activity onset, with grey representing darkness as in (E), n=8-12. (G) Time to 50% of total phase shift (PS_50_) in mice from (E), n=8-12. (H) Percent of running performed during the dark phase and (I) total distance run in 24 hours (averaged over two 24h periods), n=4-9. All analyses are mixed models with Sidak *post hoc* comparison unless otherwise noted. All data plotted as mean ± SEM.

### 5xFAD mice have accelerated circadian re-entrainment

3xTg mice carry mutations driving pathological Aβ and tau expression in the brain. To probe whether both of these pathological proteins are necessary in order to produce the re-entrainment phenotype we observed in 3xTg mice, we studied re-entrainment in 5xFAD mice. The 5xFAD model expresses mutant APP and PS1 transgenes, but no mutant tau transgene, and thus develops aggressive amyloid pathology without neurofibrillary tangles. 5xFAD mice show altered molecular circadian rhythms and circadian behavior (Lee et al., 2020; Song et al., 2015). Based on the observed behavioral phenotype in 8-month-old 3xTg mice, which have amyloid pathology but little tauopathy, we hypothesized that Aβ is sufficient to alter re-entrainment. We studied female 5xFAD mice at 7 months of age, at which time they have extensive Aβ plaques (Fig. 3A), though none are detected in the SCN (Supplementary Fig. 1C). We repeated the jet lag experiment described above in these aged 5xFAD mice (Fig. 3B). Behavior onset was significantly earlier in 5xFAD mice than WT mice on days 2 (p<.014), 3 (p<.004), and 4 (p<.002) after the shift (Fig. 3C), and mean PS_50_ was reached 2.27 days earlier (p<.006) (Fig. 3D). Free-running period and preference for running during the dark phase were not significantly affected by genotype (Fig. 3E-F).

**Figure 3:**
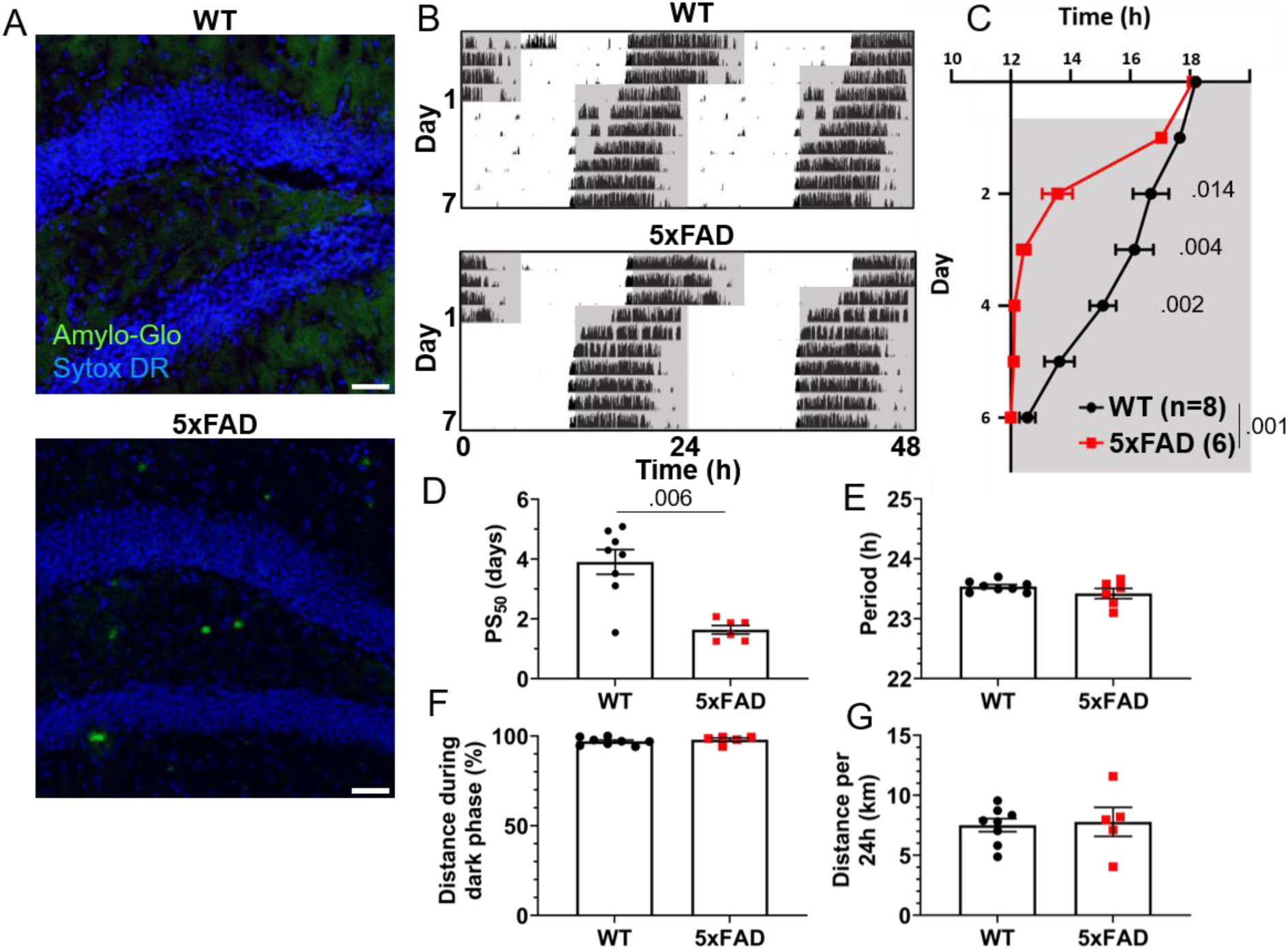
Altered circadian re-entrainment in 5xFAD mice. (A) Representative images from the hippocampus of 7mo 5xFAD and littermate control WT mice. Aβ plaques are stained with Amylo-Glo (green) and nuclei are stained with Sytox-DR (blue). Scale bars = 500um. (B) Representative double-plotted actograms of 7mo WT and 5xFAD mice subjected to a 6h phase advance. Light and dark phases of the LD cycle are represented by white and grey background, respectively. (C) Group analysis of activity onset, with grey representing darkness as in (B). Mixed model with Sidak post hoc comparison, n=6-8. (D) Time to 50% of total phase shift (PS_50_) in mice from (C), n=6-8. (E) Free-running period (averaged over 7 days) in mice maintained in constant darkness, n=6-8. (F) Percent of running performed during the dark phase and (G) total distance run in 24 hours (averaged over two 24h periods), n=6-8. All analyses are two tailed Student’s t-tests unless otherwise noted. All data plotted as mean ± SEM.

Total running distance was not altered in 5xFAD mice, suggesting that hyperactivity or perseverative behavior were not responsible for altered performance on the running wheel in this model (Fig. 3G). These results closely recapitulate the findings in aged 3xTg mice. Thus, amyloid pathology in the absence of mutant tau is sufficient to alter circadian re-entrainment in these AD mouse models.

5xFAD mice do not display the same increased bodyweight phenotype as 3xTg mice, and we again found no consistent correlation between body weight or weight loss during the running wheel period and speed of re-entrainment (Supplementary Fig. 2G-I).

To test whether abnormal tau can drive this re-entrainment phenotype in the absence of Aβ pathology, we repeated the jet lag experiment with the PS19 mouse model, which carries a mutation driving aggressive tauopathy, but does not carry amyloidogenic mutations (Yoshiyama et al., 2007). Unlike the 3xTg and 5xFAD models, re-entrainment was not altered in female PS19 mice at 7 months (Supplementary Fig. 3). Thus, the accelerated re-entrainment phenotype appears to be driven by amyloid, not tau, pathology.

### AD model mice exhibit heightened sensitivity to photic cues

Light entering via the retina entrains the SCN circadian clock. Amyloid and tau pathology are detectable in the retina in AD (Hart et al., 2016; Koronyo et al., 2017), and the cells which provide photic inputs from the retina to the circadian system, the intrinsically photosensitive retinal ganglion cells (ipRGCs), are decreased in AD (la Morgia et al., 2016). We next examined whether altered re-entrainment in 3xTg mice could be influenced by altered light sensing in the retina, rather than altered circadian timekeeping in the SCN.

We first analyzed the retinas of aged 3xTg mice by immunohistochemistry. Though we detected Aβ and tau phosphorylation in the retinas of 13-month-old 3xTg mice (Fig. 4A-B), we did not observe gross changes to retinal morphology or the RGC population (Fig. 4C), and there was no significant decrease in RGC density in 3xTg retinas (p>.298) (Fig. 4D).

**Figure 4:**
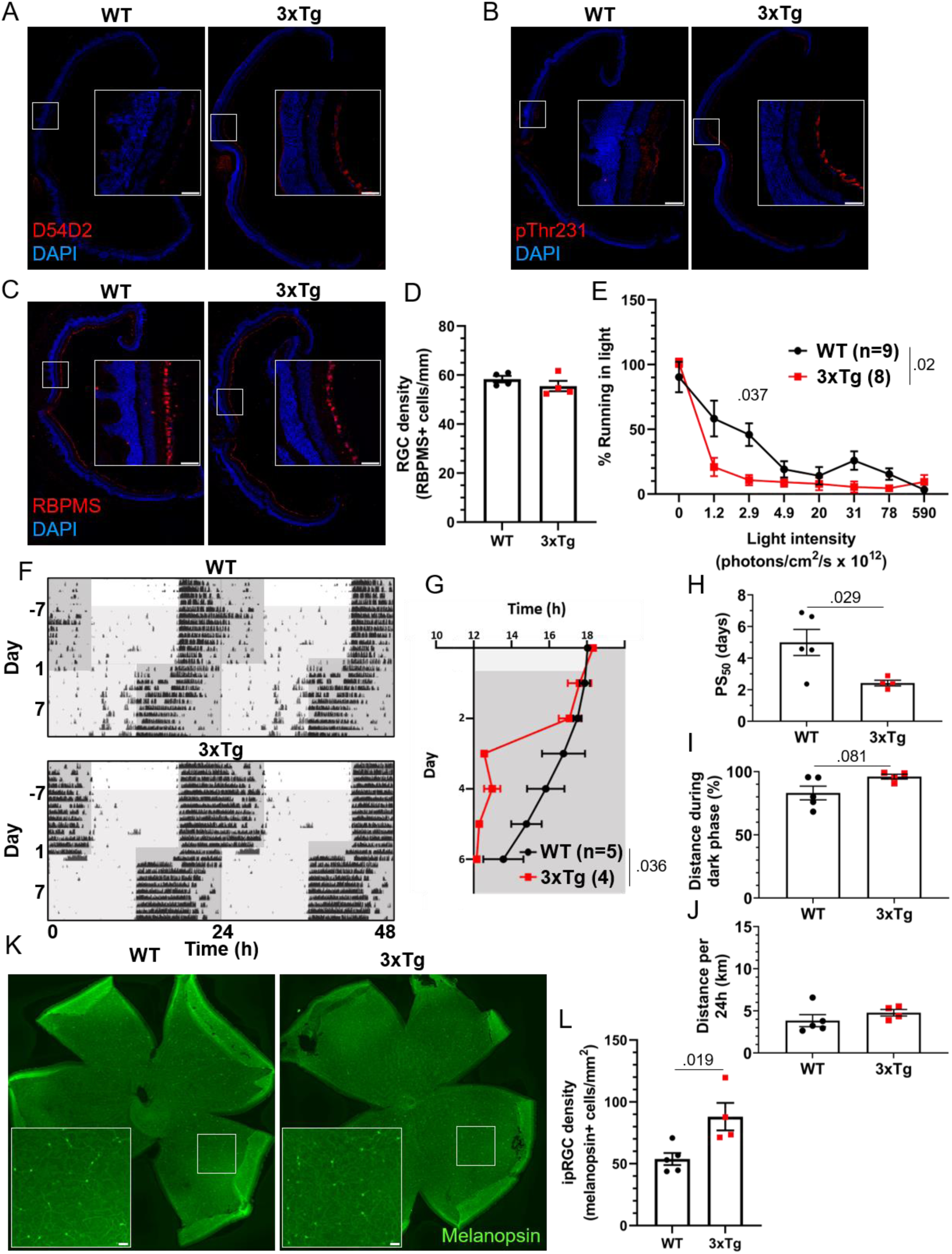
Visual circadian function is altered in 3xTg mice. 13 month old WT and 3xTg sagitally sectioned retinas were stained with DAPI and (A) D54D2 for Aβ, (B) pThr231 for phospho-tau, and (C) RBPMS for retinal ganglion cells. (D) RGC density is quantified. (E) Reduction in running caused by light pulses of increasing intensity during the dark phase in 8-month-old 3xTg and B6129SF2/J WT mice, n=8-9. Lights of different intensities were turned on from ZT13-14 and distance run in that period was compared to running during the same period of constant darkness the previous night. (F) Dim light jet lag trial representative double-plotted actograms. Lighting was switched from 590×10^12^ to 2.9×10^12^ photons/cm^2^/s. 7 days later, LD phase was advanced by 6h. Dark phase represented by dark grey background, dim light represented by light grey. (G) Group analysis of activity onset, with grey representing darkness as in (F). Mixed model with Sidak post hoc comparison, n=4-5. (H) Time to 50% of total phase shift (PS_50_) in mice from (G), n=4-5. (I) Percent of running performed during the dark phase and (J) total distance run in 24 hours (averaged over two 24h periods), n=4-5. (K) Retina whole mounts from WT and 3xTg mice stained for melanopsin to identify ipRGCs. (L) Quantification of (K). All analyses are two tailed Student’s t-tests unless otherwise noted. All data plotted as mean ± SEM. Scale bars = 50um.

The jet lag re-entrainment paradigm relies on the retina to detect photic entrainment cues and the SCN to shift the biological clock in response to those cues. We next tested negative masking, a test of behavioral response to light that depends on the retina but not the SCN. Masking is a phenomenon wherein changes in light conditions can alter normally circadian-controlled behaviors without first altering the circadian pacemaker. For example, a mouse running in the dark may stop running if the lights are suddenly turned on, in spite of it still being the animal’s active phase. To measure negative masking, we gave a one-hour pulse of light of different intensities beginning one hour after the onset of the dark phase of a 12:12 LD cycle and monitored running wheel activity of 8-month-old 3xTg and WT mice during that time. By comparing to running during uninterrupted darkness, we could calculate the degree to which different intensities of light masked running behavior, and therefore measure the sensitivity of the mouse to circadian light cues in a non-SCN-dependent system. We found a significant difference in negative masking behavior between 3xTg and WT mice (p<.02), and 3xTg running was significantly more suppressed by low-intensity 2.9×10^12^ photons/cm^2^/s lighting than WT (p<.037) (Fig. 3E). Unexpectedly, this demonstrates an elevated responsiveness to photic circadian signals in AD model mice, raising the possibility that increased perceived intensity of the photic re-entrainment cue in the jet lag paradigm may contribute to more rapid re-entrainment.

To address this question, we repeated the jet lag experiment with these mice under dim lights. Mice were switched from the 590×10^12^ photons/cm^2^/s lighting conditions used for previous experiments to only 2.9×10^12^ photons/cm^2^/s, the intensity in the masking experiment where we found the most significant difference between 3xTg and WT. Mice were kept on a 12:12 LD cycle with these dim lights for 7 days, during which time all mice maintained their entrainment (Fig. 4F). After 7 days the jet lag phase advancement was performed, still under dim lights, and re-entrainment was observed. 3xTg mice re-entrained significantly more rapidly than WT (p<.036) (Fig. 4G), with a PS_50_ 2.54 days earlier (p<.029) (Fig. 4H). This was a more dramatic difference between groups than was observed in either 8 or 13 month old mice under bright lights (Fig. 1D, J). While dim light PS_50_ for 3xTg mice was slightly higher than those in bright lights (PS_50_ = 2.42 in dim light, 1.68 in bright light at 8 months, 2.16 in bright light at 13 months), it was dramatically higher in WT mice under dim lights compared to under bright lights (PS_50_ = 5.00 in dim light, 2.74 in bright light at 8 months, 3.57 in bright light at 13 months). Also notably, after re-entrainment under dim light, some WT animals displayed unusual running behavior including considerable bouts of running before the onset of the dark phase (see Fig. 4F, upper panel). This was not observed in 3xTg mice. This resulted in a trend (p=.081) towards decreased preference for running in the dark phase in WT mice (Fig. 4I). Total running was not significantly different between the groups (Fig. 4J). Overall, the masking and dim light experiments demonstrated that 3xTg mice respond very similarly to dim light as to bright light while WT mice are much less responsive to dim light, both as a masking stimulus and an entrainment cue. This supports the hypothesis that altered retinal function contributes to differing circadian behavior. We also found a 64% increase in the density of melanopsin-positive ipRGCs in the retinas of these 3xTg mice compared with WT (p<.019). All retinal tissue samples were collected between zeitgeber time (ZT) 5-8 and ipRGC density was averaged across images from multiple regions of the retina to minimize the effects of rhythmicity(Hannibal et al., 2005) and regional disparities(Dacey et al., 2005) in melanopsin expression. This increase in the ipRGC population may contribute to the heightened sensitivity to light as a circadian cue in 3xTg mice.

## Discussion

We found that two amyloid-based AD mouse models have significantly accelerated re-entrainment in a jet lag paradigm. This behavioral phenotype was not affected by the depletion of microglia and did not depend on the presence of tauopathy. These mice were also more sensitive to light in a trial of negative masking, an SCN-independent behavior, and had increased density of ipRGCs. Together these results uncover a novel circadian phenotype in AD model mice driven by Aβ.

This is the first report, to our knowledge, of altered jet lag behavior in AD model mice. Previous studies in APP/PS1 mice (Kent et al., 2019; Otalora et al., 2012) and male 3xTg mice (González-Luna et al., 2021) did not show altered re-entrainment in the jet lag paradigm. However, the APP/PS1 model has been criticized as not replicating some circadian phenotypes observed in AD patients (Sheehan & Musiek, 2020) and most studies find less severe pathophysiology and behavioral phenotypes in male than female 3xTg mice (Dennison et al., 2021). Given the variability between different AD models, our finding of the same robust re-entrainment phenotype in two different models is important for validating that this phenotype is not the result of peculiarities of a specific genetic model. We do not observe alterations to free running period or daily activity patterns, two measures of circadian behavior in which differences have been reported in some studies with 3xTg mice (Adler et al., 2019; Wu et al., 2018). Previous research has not found altered free running period in 5xFAD mice (Nagare et al., 2020), consistent with our findings here. We did observe notable differences between WT mice in our studies depending on age (Fig. 1D, J) and strain (Fig. 1D, Fig. 3D, Supplementary Fig. 3C). Indeed, given the very rapid re-entrainment of littermate controls of PS19 mice, the B6C3 background strain in this model may not be well suited to detecting fine differences in re-entrainment behavior, and other tauopathy models may exhibit behavioral differences. These heterogeneous results highlight the importance of sex, age, and background strain in circadian behavioral experiments.

Modulating neuroinflammation in AD is a promising field of research, and microglia can both contribute to and protect against disease progression. As neuroimmune activation and circadian disruptions both appear early in the progression of AD, and inflammation can modulate circadian rhythms, microglia seemed a promising possible mechanism underlying circadian symptoms. However, we found that microglia depletion did not rescue jet lag behavior in 3xTg mice. The microglia depletion in our experiment was acute and longer treatment may have other effects on circadian behavior, but our results suggest that acutely targeting inflammation in AD may not directly ameliorate circadian disruptions. Retinal microglia are also likely not acutely involved in the jet lag phenotype as CSF1R inhibitors effectively deplete microglia in the retina as well as in other regions of the brain (Dharmarajan et al., 2017; Huang et al., 2018). We did observe a trend towards accelerated re-entrainment in WT mice after microglia depletion, which may suggest a role for microglia in circadian regulation in the healthy brain. A study reported altered circadian behavior in rats after using a targeted diphtheria toxin approach to deplete microglia (Sominsky et al., 2021), although these results may have been influenced by sickness behavior induced by the diphtheria toxin approach. Another recent study found that microglia depletion alters sleep in mice (H. Liu et al., 2021), but did not study circadian-regulated behaviors. More research is needed to understand the effects of microglia on circadian rhythms in health and disease.

Accelerated recovery from jet lag is not one of the circadian symptoms of AD, but this phenotype may reflect underlying circadian disruptions. Strategies for managing sundowning and improving sleep-related symptoms in AD patients include maintaining strict light schedules, increasing exposure to intense light during daylight hours, and decreasing exposure to light during the night (Mitolo et al., 2018). AD patients also have decreased amplitude of rhythmic circadian gene expression (Cermakian et al., 2011). This may indicate a weak biological circadian clock, making them heavily dependent on external cues for its maintenance. Rapid re-entrainment may indicate a decrease in the coupling and synchrony of neurons in the SCN, resulting in an intrinsic circadian timekeeper less resistant to being shifted by misaligned photic entrainment cues. A very similar jet lag phenotype is observed in mice genetically knocked out for vasopressin signaling (Yamaguchi et al., 2013), which impairs interneuronal communication in the SCN. Some reports find decreased AVP expression in the SCN of AD patients (Harper et al., 2008; R. Y. Liu et al., 2000), though this has been disputed (Wang et al., 2015), and previous reports have found decreased expression of AVP in the SCN of 3xTg mice (Sterniczuk et al., 2010). Further studies will be needed to determine if changes in SCN signaling also contribute to this phenotype and how amyloid pathology drives those changes.

Our finding of increased sensitivity to low-intensity photic circadian cues in an AD model suggests mechanisms outside the SCN may play a role in the jet lag phenotype. A previous study found increased negative masking in male 3xTg mice (González-Luna et al., 2021) but did not examine behavior at low light levels. The retinal degeneration in AD models, including 3xTg mice (Grimaldi et al., 2018), and ipRGC loss in AD (la Morgia et al., 2016) suggest decreased sensitivity, not increased. The limited research on visual ability in 3xTg mice has not reported deficits (King et al., 2018), while some decrease in RGC activity can be observed by electroretinography (Frame et al., 2022). Recent research found a loss of ipRGCs and decreased ipRGC projections to the SCN in the APP/PS1 model (Carrero et al., 2023). Our finding in 3xTg mice of heightened responses to photic cues in two circadian behavioral paradigms, as well as increased ipRGC density, runs contrary to these findings and suggests that in some models, or at some stages of disease progression, ipRGC signaling may be increased rather than decreased. More research will be necessary to determine the causes of these differences between models or disease stages. The heterogeneity of the ipRGC population should also be further studied in these mice. Some ipRGC subtypes express melanopsin at very low levels and would be undetectable by the IHC technique used here. One possible explanation for the increase in melanopsin+ cells in the 3xTg retinas could be an increase in melanopsin expression in non-M1 ipRGCs, or a skew towards more M1 and M2 ipRGCs, rather than an increase in the number of the ipRGC population. As these non-M1 ipRGCs have different functions and projection patterns than M1 ipRGCs (reviewed in (Do, 2019)), such a change in the ipRGC population could have consequences for diverse visual circuits and behaviors (Panda et al., 2003).

In summary, we demonstrate that AD model mice exhibit strikingly altered circadian behavior, suggesting a heightened sensitivity to photic circadian cues. This appears to be driven by pathogenic Aβ and does not require the presence of microglia or mutant tau.

## Methods

### Mice

All animal experiments were conducted in accordance with the University of Virginia Institutional Animal Care and Use Committee. Animals were housed in a temperature and humidity controlled vivarium (22- 24°C, ∼40% humidity) and were provided with food and water ad libitum. 3xTg experiments were conducted with 8-13 month old female 3xTg mice on a B6129 background (Oddo et al., 2003) (Jackson Laboratory #034830), with age-matched B6129SF2/J (Jackson Laboratory #101045) females as wild type controls. 5xFAD experiments were conducted with 7 month old female heterozygous 5xFAD mice on a C57BL/6J background (Oakley et al., 2006) (Jackson Laboratory #034848), with littermates genotyped as not expressing the mutant transgene serving as wild type controls. PS19 experiments were conducted with 7 month old female heterozygous PS19 mice on a B6C3 background (Yoshiyama et al., 2007) (Jackson Laboratory #008169), with littermates genotyped as not expressing the mutant transgene serving as wild type controls. For microglia depletion experiments, mice were given chow formulated with PLX3397 (660mg/kg) or control chow for 7 days before light cycle shift and were maintained on PLX or control chow for the remainder of the experiment.

### Behavioral analysis

Behavioral testing protocol was adapted from (Grippo et al., 2017). Mice were individually housed in cages (Nalgene) containing running wheels in light-tight boxes which were illuminated with timed fluorescent lights (590×10^12^ photons/cm^2^/s). Wheel running data were collected and analyzed with ClockLab software (Actimetrics). Activity onset was automatically detected by ClockLab software and when necessary, corrected by eye by an experimenter blinded to genotype and treatment group. Mice were allowed to habituate to running wheel cages and entrained to a 12hr:12hr LD cycle for at least 7 days before experiments began.

In jet lag re-entrainment trials, the onset of the dark phase was abruptly advanced by 6 hours and running wheel activity was recorded for at least 7 days after light cycle shift. PS_50_ values were calculated using Prism software (GraphPad) by fitting a sigmoid dose-response curve to onset times in days 0-6 after light cycle shift (Kiessling et al., 2010). Total running distance and preference for running in the dark were measured after all mice had completely re-entrained after a phase shift and were averaged across 2 days. To determine free running period, after all mice had completely re-entrained they were switched to DD and period was calculated from the onset of activity across 7 days.

In masking trials, light intensity was decreased by wrapping fluorescent lights in neutral density filter films and measured with a spectrometer (Sekonic Spectrometer C-800). Illuminance was measured in lux, and photon flux, summed from 380 to 780nm, was calculated using formulas in the supplementary materials in Lucas et al., 2014. A 1hr light pulse was delivered from ZT13-14. Percent running in light was calculated by comparing to the activity measured during the same period of constant darkness on the preceding day for each mouse. Individual trials were separated from each other by 3-4 days.

### Histological analysis and imaging

All mice used for histological analysis were sacrificed from ZT 5-9. Mice were anesthetized with a ketamine:xylazine solution and transcardially perfused with chilled phosphate buffered saline (PBS) followed by 4% paraformaldehyde (PFA). Brains were dissected and post-fixed in 4% PFA for 24h at 4°C. Brains were cryoprotected by incubating >24h in 15% sucrose in PBS followed by 30% sucrose before being frozen and sectioned at 30μm with a cryostat (Leica CM 1950). For stains using primary antibodies raised in mice, sections were blocked using Mouse on Mouse blocking solution (Vector Laboratories) for 2h at room temperature. Otherwise, brain sections were incubated with blocking solution (5% bovine serum albumin, 2% horse serum, 1% Triton X-100 in PBS) for 2h at room temperature. Primary antibodies were diluted in blocking solution and placed on sections overnight at 4°C. Sections were washed and incubated with secondary antibodies, diluted in blocking solution, for 2h at room temperature. Sections were washed and if applicable were treated with DAPI (Sigma-Aldrich) (1:2000 in PBS for 15min), Sytox Deep Red (Thermo Scientific) (1:2000 in PBS for 30min), and/or AmyloGlo (Biosensis) per manufacturer protocol. Sections were washed and then mounted with ProLong Gold mounting media (Thermo Fisher).

Retinas were dissected and stained as in (Gao et al., 2022) with minor modifications. For retinal sectioning, eye cups were dissected and post-fixed in 4% PFA for 30min at room temperature. Eye cups were cryprotected in 30% sucrose in PBS overnight at 4°C, frozen, and sectioned on the cryostat at 14μm. Sections were again post-fixed with 2% PFA for 30min at room temperature. Sections were then blocked, stained, and mounted as described for brain sections above. For retina whole-mounts, eye cups were dissected and post-fixed with 2% PFA for 1h on ice. They were then washed and blocked with blocking solution described above for 2h at room temperature. They were incubated with primary antibody diluted in blocking solution overnight on a shaker at 4°C. They were then washed and incubated with secondary antibody diluted in blocking solution overnight on a shaker at 4°C. Finally the retinas were removed from the eye cup, cut to create four quadrants, and flat mounted with ProLong Gold mounting media.

Primary antibodies used were anti-phospho-tau AT180 (mouse, Invitrogen MN1040, 1:250), anti-phospho-tau pThr231 (rabbit, Invitrogen 701056, 1:500), anti-RBPMS (rabbit, Abcam ab152101, 1:500), anti-Iba1 (rabbit, Wako 019-19741, 1:300), anti-amyloid D54d2 (rabbit, Cell Signaling 8243, 1:200), and anti-melanopsin (Panda et al., 2002) (rabbit, 1:2500). Secondary antibodies used were donkey anti-rabbit AlexaFluor 594 (Invitrogen A21207), donkey anti-rabbit AlexaFluor 647 (Invitrogen A32795), donkey anti-mouse AlexaFluor 594 (Invitrogen A21203), and donkey anti-mouse AlexaFluor 647 (Invitrogen A11012).

Images were acquired with a Keyence BZ-X800 fluorescence microscope and images were stitched using BZ-X800 Analyzer software. Cell quantification was performed with ImageJ. For quantification of microglia, a 0.36mm^2^ square was drawn in the ventromedial hypothalamus and all Iba1+ cells were counted. For quantification of RGCs in sectioned retinal tissue, 500-700µm lines were drawn along the RGC layer beginning 300µm to either side of the optic nerve exit point and all RBPMS+ cells were counted. For quantification of ipRGCs in retina whole mounts, four 0.25mm^2^ squares were drawn in each retina at equal distances from the optic nerve exit point and melanopsin+ cells were counted in each, with cell density being calculated as an average of these counts.

## Acknowledgements

We thank R. Grippo and Q. Tang for help with behavioral experiments, J. Cole for assistance with retinal staining, U. Eyo for providing PLX3397 chow, Virginia Alzheimer’s Disease Center for providing antibodies, and I. Provencio for advice and antibodies. This research was funded by grants from the Owens Family Foundation, Commonwealth of Virginia’s Alzheimer’s and Related Diseases Research Fund and K08DK097293 to HAF, T32GM139787 to TKW and R35 GM140854 to ADG.

