## Supplementary figures for "Altered circadian behavior and light sensing in mouse models of Alzheimer’s disease"

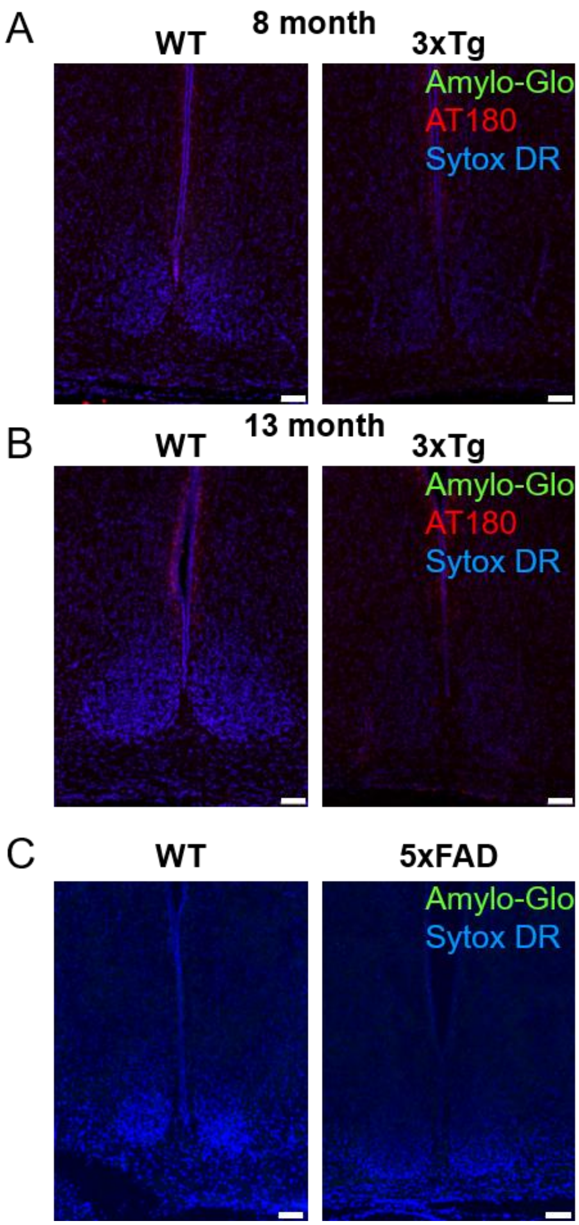


Supplementary Figure 1: Amyloid and tau pathology not detected in the SCN of AD models

Stains of the SCN and surrounding region with Sytox-DR for nuclei (blue), AmyloGlo for Aβ plaques (green), and AT180 for phosphorylated tau (red) in WT and 3xTg mice at (A) 8 months, and (B) 13 months.

(C) Stains with Sytox-DR for nuclei and AmyloGlo for Aβ plaques in WT and 5xFAD mice at 7 months.

Scale bars =100um.

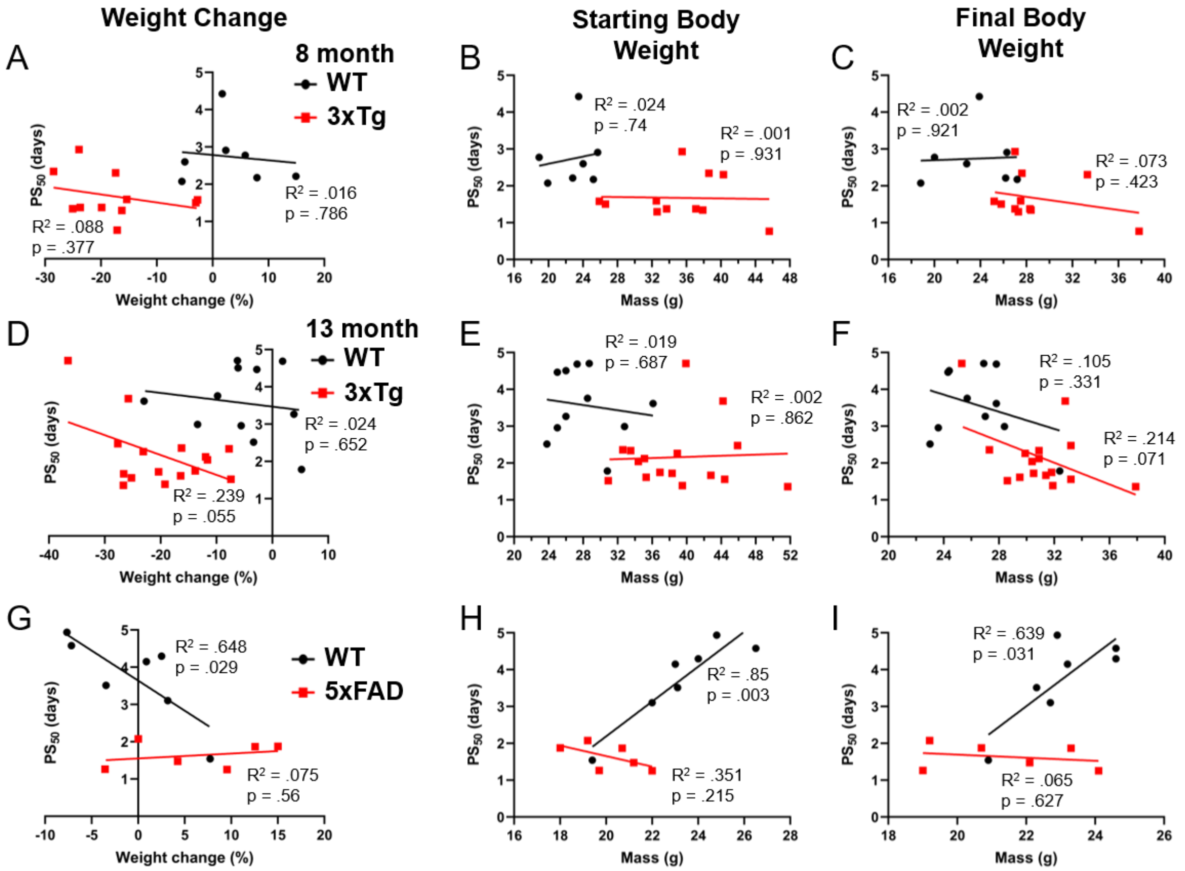


Supplementary Figure 2: Associations of weight loss and body weight with jet lag re-entrainment.

(A) Percent weight change over the course of wheel running in 8-month-old WT and 3xTg mice plotted with time to 50% re-entrainment ($PS_{50}$).

(B) Body weight at the beginning of wheel running plotted with $PS_{50}$.

(C) Body weight at the end of wheel running plotted with $PS_{50}$.

(D-F) Same as (A-C) with 13-month-old WT and 3xTg mice.

(G-I) Same as (A-C) with 7-month-old WT and 5xFAD mice.

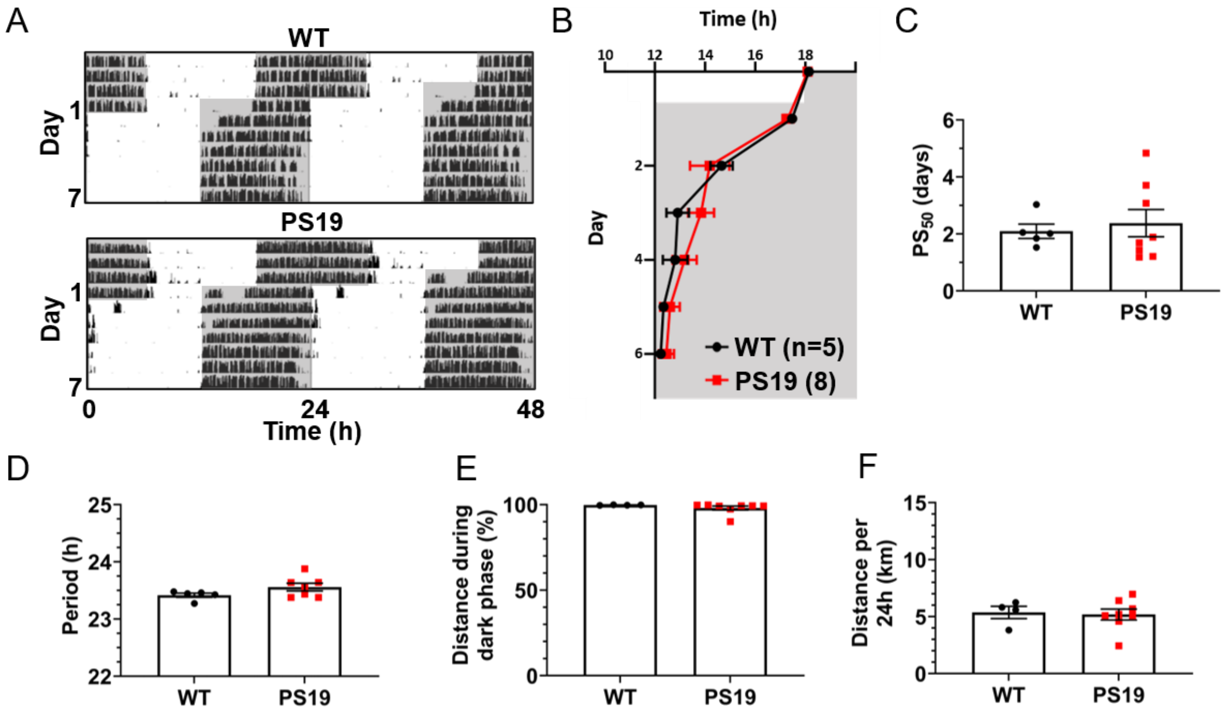


Supplementary Figure 3: No altered circadian re-entrainment in PS19 mice.

(A) Representative double-plotted actograms of 7mo PS19 and littermate control WT mice subjected to a 6h phase advance. Light and dark phases of the LD cycle are represented by white and grey backgrounds, respectively.

(B) Group analysis of activity onset, with grey representing darkness as in (A). Mixed model with Sidak post hoc comparison, n=5-8.

(C) Time to 50% of total phase shift ($PS_{50}$) in mice from (B), n=5-8.

(D) Free-running period (averaged over 7 days) in mice maintained in constant darkness, n=5-7.

(E) Percent of running performed during the dark phase and (F) total distance run in 24 hours (averaged over two 24h periods), n=4-8.

All analyses are two tailed Student's t-tests unless otherwise noted. All data plotted as mean ± SEM.
